# A luciferase-based assay for assessing IRES-mediated translation in Wheat Germ Extract

**DOI:** 10.64898/2026.04.07.716985

**Authors:** Max Cortot, Thorsten Stehlik, Aline Koch, Timo Schlemmer

## Abstract

Efficient protein synthesis in eukaryotic cells typically requires a 5′ cap structure on messenger RNAs (mRNAs). However, under stress conditions or in viral infection, translation can also occur independently of the cap via internal ribosomal entry sites (IRES). IRES elements are therefore key regulators of protein expression in both viral and cellular contexts. Here we describe a cell-free protocol to quantitatively assess IRES-mediated translation using wheat germ extract (WGE) and a firefly luciferase (FLuc) reporter. The protocol includes template preparation, RNA synthesis and luminescence measurement following *in vitro* translation in WGE. This method enables rapid and robust comparison of IRES activity under controlled conditions and can additionally be applied to evaluate mRNA modifications designed to enhance translation efficiency.

**Key features:**

- Stringent *in vitro* workflow from DNA template preparation through RNA synthesis and protein synthesis to reporter readout, including quality controls.
- Evaluation of IRES-driven translation suitable for testing combinations of IRES and CDS.
- translation analysis <u>without</u> radioactive labeling.

**Graphical overview:** 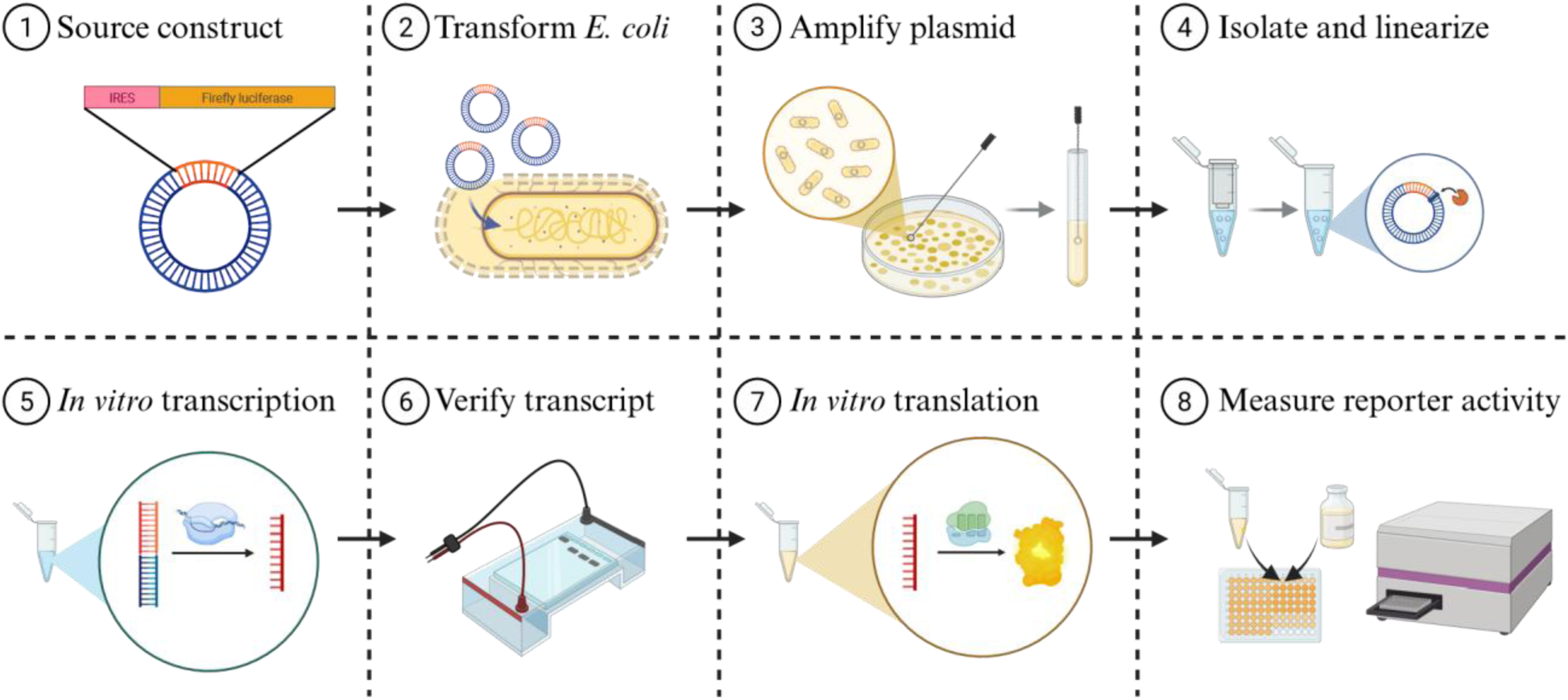

**Graphical Abstract:** **Pipeline for the production and evaluation of IRES-firefly luciferase constructs using wheat germ extract. (1-4) Preparation:** IRES-firefly luciferase constructs are amplified in *E. coli* and isolated from bacterial cells. Plasmids are linearized to prepare for *in vitro* transcription. **(5-6) Transcript synthesis and verification:** In vitro transcription is followed by electrophoretic validation to confirm integrity and correct molecular weight. **(7-8) Translation and detection:** Translation is executed in wheat germ extract and quantified by measuring reporter activity in a luminometer.

## Background

Viruses have evolved several strategies to harness the host translational machinery. One such strategy is the development of internal ribosomal entry sites (IRES) typically located in the 5’ untranslated region (5’ UTR) of a viral mRNA. IRES are cis-acting RNA elements with complex secondary structures that enable cap-independent translation initiation [1]. While in hosts cap-dependent translation is more efficient under normal conditions, IRES-mediated translation becomes increasingly relevant during cellular stress [2], allowing some cellular stress response mRNAs containing IRES sequences, such as maize heat shock protein 101, cap-independent translation during heat stress [3,4]. Beyond natural stress responses, IRES sequences are essential for the translation of circular RNAs, which lack free ends required for canonical, cap-dependent translation [5]. Because IRES activity can vary depending on their viral or cellular origin and the host organism, evaluating different IRES sequences is critical for optimizing translation efficiency in biotechnological applications [6]. Wheat germ extract (WGE) is a cell-free system for protein synthesis. WGE possesses eukaryotic translation machinery and can translate a variety of eukaryotic transcripts with high yield and does not require codon optimization [7]. Another advantage is the very high solubility and bioactivity of the expressed proteins, increasing the amount of functional proteins for downstream applications [8]. Computational predictions showed a 90.9% solubility rate of the human proteome in the WGE system, compared to 35.5% in *E. coli* expression systems [9]. The cell-free environments allows control of the experimental conditions through additives such as surfactants, detergents, peptides or amphipols [8]. This, combined with the easily quantifiable reporter outputs, makes WGE an ideal system to compare relative IRES activity.

Firefly luciferase (FLuc) is routinely used as reporter in biological assays. The luciferase gene, which originates from the firefly *Photinus pyralis,* was first cloned and expressed in mammalian cells in the late 80s [10,11]. The enzyme has a molecular weight of 62 kDa and requires D-luciferin, ATP, oxygen and a metal cation (such as Mg^2+^) as substrates [12]. Luciferase enzymes have the advantage of illuminating the dynamic changes in reporter transcription, contrary to fluorescent protein reporters which have longer intracellular protein half-lives [12,13]. In the case of the endpoint measurement described in this protocol, ease of detection and the large linear dynamic range are the key advantages of the FLuc reporter

Here we provide a step-by-step protocol starting from generation of IRES-containing plasmids, followed by plasmid purification and linearization for *in vitro* transcription, *in vitro* translation in wheat germ extract, and quantification of IRES activity using a firefly luciferase reporter (Fig. 1). We provide guidance for control experiments at each step for self-control for future users. Taken together, this protocol is the first one dealing with IRES evaluation for relative quantification of different IRES-reporters and allows rapid setting up of *in vitro* translation and activity measurements of IRES sequences. Our plasmid set functions as a positive control and resource for testing combinations of plant viral IRES elements in plants.

**Figure 1.**
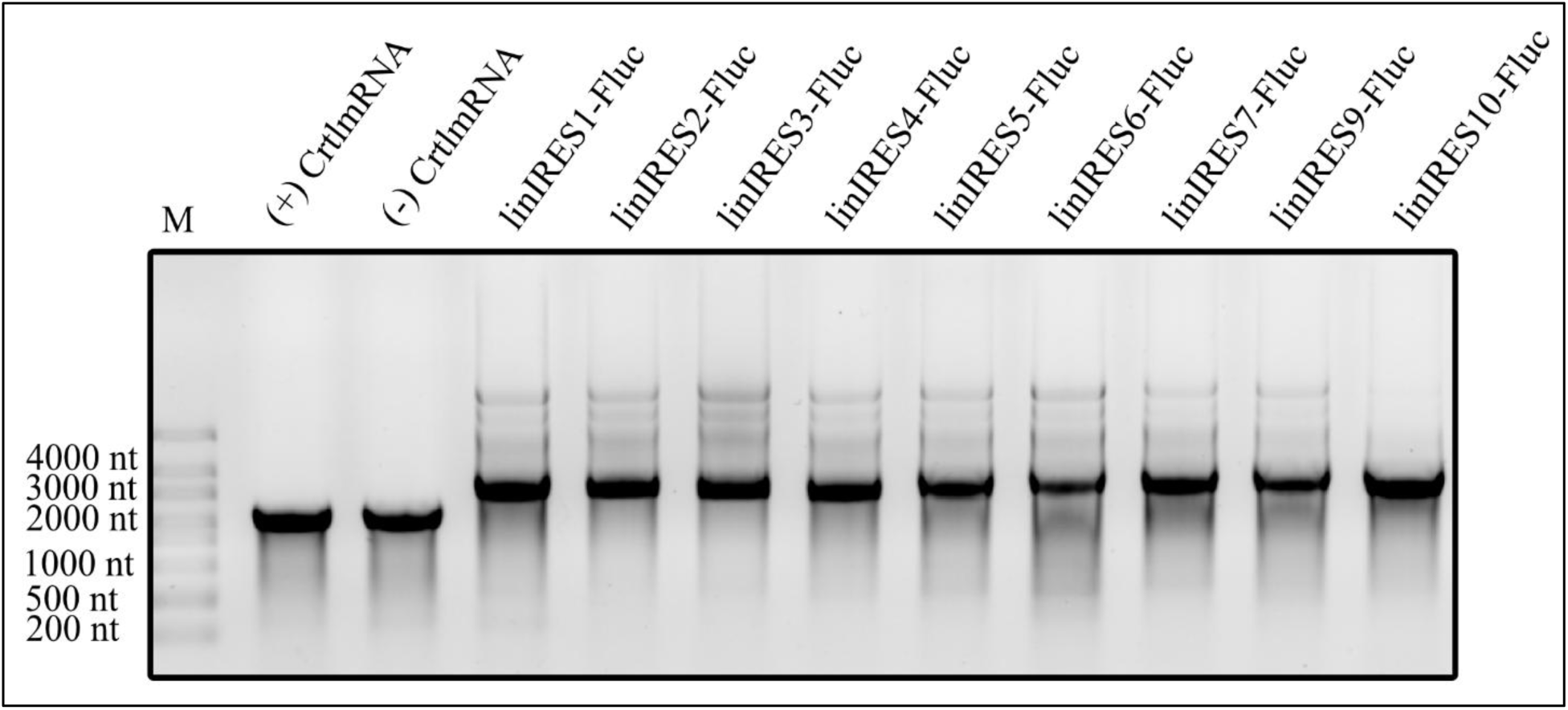
Gel electrophoresis analysis of different transcripts. All RNAs migrate at their expected sizes, as detailed in Table 22. The (+) CtrlmRNA and (-) CtrlmRNA lanes correspond to the RNA transcribed from the L4821 plasmid, with (+) possessing a 3′-O-Me-m^7^G(5’)ppp(5’)G RNA Cap Structure Analog and (-) possessing no cap structure. 1.5 µL RiboRuler High Range RNA Ladder (catalog number: SM1821) were investigated as size standard (M).

## Materials and reagents

### Biological materials

1. Luciferase T7 Control DNA (Promega, catalog number: L4821)
2. NEB^®^ Stable Competent *E. coli* (High Efficiency) (New England Biolabs, catalog number: C3040)
3. pUC19mut_circIRES1_FLuc_3HA (under submission)
4. pUC19mut_circIRES2_FLuc_3HA (under submission)
5. pUC19mut_circIRES3_FLuc_3HA (under submission)
6. pUC19mut_circIRES4_FLuc_3HA (under submission)
7. pUC19mut_circIRES5_FLuc_3HA (under submission)
8. pUC19mut_circIRES6_FLuc_3HA (under submission)
9. pUC19mut_circIRES7_FLuc_3HA (under submission)
10. pUC19mut_circIRES9_FLuc_3HA (under submission)
11. pUC19mut_circIRES10_FLuc_3HA (under submission)

**Note:** Additional information on IRES origins is found on addgene. IRES denomination within plasmids was based on a listing after literature research and for eased laboratory use. The denomination is irrelevant for conducting the following protocol. Cloning of IRES8 was aborted during the research project.

## Reagents

1. 1 kb Plus DNA Ladder (New England Biolabs, catalog number: N3200)
2. 2-Propanol (Thermo Scientific, CAS number: 67-63-0)
3. 3′-O-Me-m^7^G(5’)ppp(5’)G RNA Cap Structure Analog (New England Biolabs, catalog number: S1411S)
4. AfeI (New England Biolabs, catalog number: R0652S)
5. Agar-Agar, Kobe I (Carl ROTH, CAS number: 9002-18-0)
6. Agarose NEEO Ultra-Quality (Carl ROTH, catalog number: 2267.4)
7. Amino Acid Mixture, complete (Promega, catalog number: L4461)
8. Ammonium peroxydisulphate (Carl ROTH, catalog number: 9592.2)
9. Ampicillin (Sigma Aldrich, CAS number: 69-52-3)
10. BIS-TRIS (Carl ROTH, catalog number: 9140.3)
11. Boric acid (Carl ROTH, catalog number: 6943.6)
12. Bovine Serum Albumin (BSA) (Carl ROTH, catalog number: 3737.1)
13. Bromophenol blue (Carl ROTH, catalog number: T116.3)
14. Clarity Western ECL Substrate (Bio-Rad, catalog number: 1705061)
15. DNase I (New England Biolabs, catalog number: M0303S)
16. DraIII-HF^®^ (New England Biolabs, catalog number: R3510)
17. EDTA, 0.5 M Solution (Thermo Fisher Scientific, catalog number: J15694.AP)
18. EDTA, sodium salt (Carl ROTH, catalog number: 8043.2)
19. Ethanol, ≥99,8 %, p.a. (Carl ROTH, catalog number: 9065.4)
20. Formamide deionized (Carl ROTH, catalog number: P040.1)
21. Glacial acetic acid (Merck, catalog number: 33209)
22. Glycerol Formal (Carl ROTH, catalog number: 0798.3)
23. Glycine (Merck, catalog number: 33226)
24. HA Tag Recombinant monoclonal antibody (Proteintech, catalog number: 81290-1-RR)
25. HiScribe^®^ T7 Quick High Yield RNA Synthesis Kit (New England Biolabs, catalog number: E2050S)
26. Lithium Chloride (Carl ROTH, CAS number: 7447-41-8)
27. Methanol (Thermo Fisher Scientific, CAS number: 67-56-1)
28. Monarch^®^ Plasmid Miniprep Kit (New England Biolabs, catalog number: T1010)
29. Nuclease-free water, prepared using a Milli-Q^®^ Advantage A10 with a Biopak^®^ Endfilter (Merck Millipore, Z00Q0V0T0 and CDUFBI001)
30. ONE-Glo™ Luciferase Assay System (Promega, catalog number: E6110)
31. Peroxidase IgG Fraction Monoclonal Mouse Anti-Rabbit IgG, light chain specific (Jackson Immuno Research, catalog number: 211-032-171)
32. PIPES (Carl ROTH, catalog number: 9156.3)
33. Ponceau S (Carl ROTH, catalog number: 5938.1)
34. Potassium Acetate (Merck, CAS number: 127-08-2)
35. Powdered milk (Carl ROTH, catalog number: T145.3)
36. Precision Plus Protein™ Dual Xtra Prestained Protein Standards (Bio-Rad, catalog number: 1610377)
37. Qubit^™^ RNA Broad Range Assay Kit (Thermo Fisher Scientific, catalog number: Q10210)
38. rCutsmart^™^ Buffer (New England Biolabs, catalog number: B6004S)
39. RiboRuler High Range RNA Ladder (Thermo Fisher Scientific, catalog number: SM1821)
40. RiboRuler Low Range RNA Ladder (Thermo Fisher Scientific, catalog number: SM1831)
41. RNase Inhibitor, Murine (New England Biolabs, catalog number: M0314)
42. ROTI^®^GelStain (Carl ROTH, catalog number: 3865.1)
43. ROTIPHORESE^®^ Gel 40 (37.5:1) (Carl ROTH, catalog number: T802.1)
44. SDS ultra-pure (Carl ROTH, catalog number: 2326.2)
45. Sodium acetate (Carl ROTH, CAS number: 127-09-3)
46. Sodium chloride (Carl ROTH, catalog number: 0601.3)
47. TEMED (Carl ROTH, catalog number: 2367.3)
48. TRIS (Carl ROTH, catalog number: 4855.2)
49. TRIS hydrochloride (Carl ROTH, catalog number 9090.4)
50. Tryptone (Carl ROTH, catalog number: 8952.2)
51. Tween^®^ 20 (Sigma-Aldrich, catalog number: P1379-100ML)
52. Wheat Germ Extract (Promega, catalog number: L418A)
53. Xylene cyanole (Carl ROTH, catalog number: A513.1)
54. Yeast Extract (Carl ROTH, catalog number: 2363.2)

## Solutions

(listed as required during procedure)

1. LB medium (Table 1)
2. 10x TBE buffer (Table 2)
3. TRIS hydrochloride 1 M stock solution (pH 6.8) (Table 3)
4. 6x DNA loading dye (Table 4)
5. Sodium acetate solution (3 M) (Table 5)
6. Lithium chloride solution (8 M) (Table 6)
7. Ethanol 70% [v/v] (store at -20°C) (Table 7)
8. 10x BPTE buffer (Table 8)
9. 2x RNA loading dye (Table 9)
10. TRIS hydrochloride 500 mM stock solution (pH 6.8) (Table 10)
11. TRIS hydrochloride 1.5 M stock solution (pH 8.8) (Table 11)
12. 10% SDS stock solution (Table 12)
13. 2x SDS-PAGE sample loading buffer (Table 13)
14. 10% Ammonium peroxydisulphate stock solution (Table 14)
15. Transfer buffer (semi-dry) (Table 15)
16. Ponceau S staining solution (Table 16)
17. 10x TBS buffer (Table 17)
18. TBS-T buffer (Table 18)
19. Blocking buffer (Table 19)

**Table 1.** LB medium. Aliquot and autoclave immediately.

| Reagent | Final concentration | Quantity or Volume |
| --- | --- | --- |
| Yeast Extract | 10 g/L | 50 g |
| Tryptone | 5 g/L | 25 g |
| Sodium chloride | 10 g/L | 50 g |
| Nuclease-free water | n/a | Fill to 5000 mL |
| Total (optional) | n/a | 5000 mL |

**Table 2.**
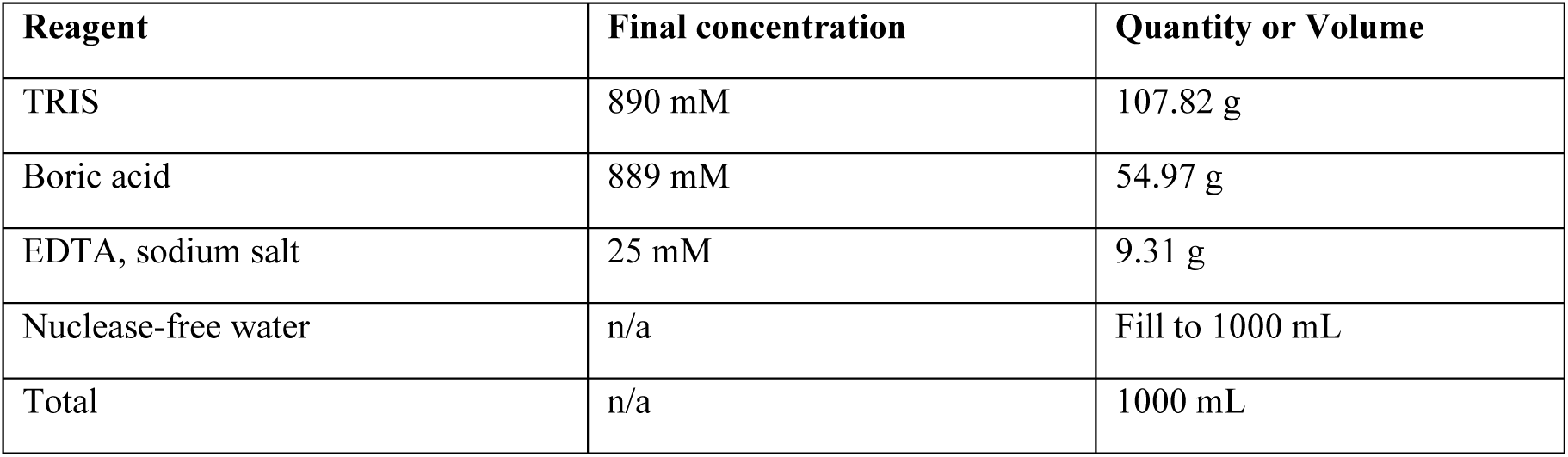
10x TBE buffer.

**Table 3.** TRIS hydrochloride 1M stock solution (pH 6.8). Adjust pH to 6.8.

| Reagent | Final concentration | Quantity or Volume |
| --- | --- | --- |
| TRIS hydrochloride | 1 M | 157.6 g |
| Nuclease-free water | n/a | Fill to 1000 mL |
| Total | n/a | 1000 mL |

**Table 4.** 6x DNA loading dye.

| Reagent | Final concentration | Quantity or Volume |
| --- | --- | --- |
| TRIS hydrochloride 1 M stock solution | 10 mM | 10 mL |
| Glycerol Formal | 60% [v/v] | 6 mL |
| Bromophenol blue | 0.03% [w/v] | 3 mg |
| Xylene cyanole | 0.03% [w/v] | 3 mg |
| Nuclease-free water | n/a | Fill to 10 mL |
| Total | n/a | 10 mL |

**Table 5.** Sodium acetate solution (3 M)

| Reagent | Final concentration | Quantity or Volume |
| --- | --- | --- |
| Sodium acetate | 3 M | 24.6 g |
| Nuclease-free water | n/a | Fill to 100 mL |
| Total (optional) | n/a | 100 mL |

**Table 6.** Lithium chloride solution (8 M).

| Reagent | Final concentration | Quantity or Volume |
| --- | --- | --- |
| Lithium chloride | 8 M | 33.92 g |
| Nuclease-free water | n/a | 100 mL |
| Total (optional) | n/a | 100 mL |

**Table 7.** 70% Ethanol.

| Reagent | Final concentration | Quantity or Volume |
| --- | --- | --- |
| Ethanol, ≥99,8 %, p.a. | 70% [v/v] | 70 mL |
| Nuclease-free water | 30% [v/v] | 30 mL |
| Total (optional) | n/a | 100 mL |

**Table 8.**
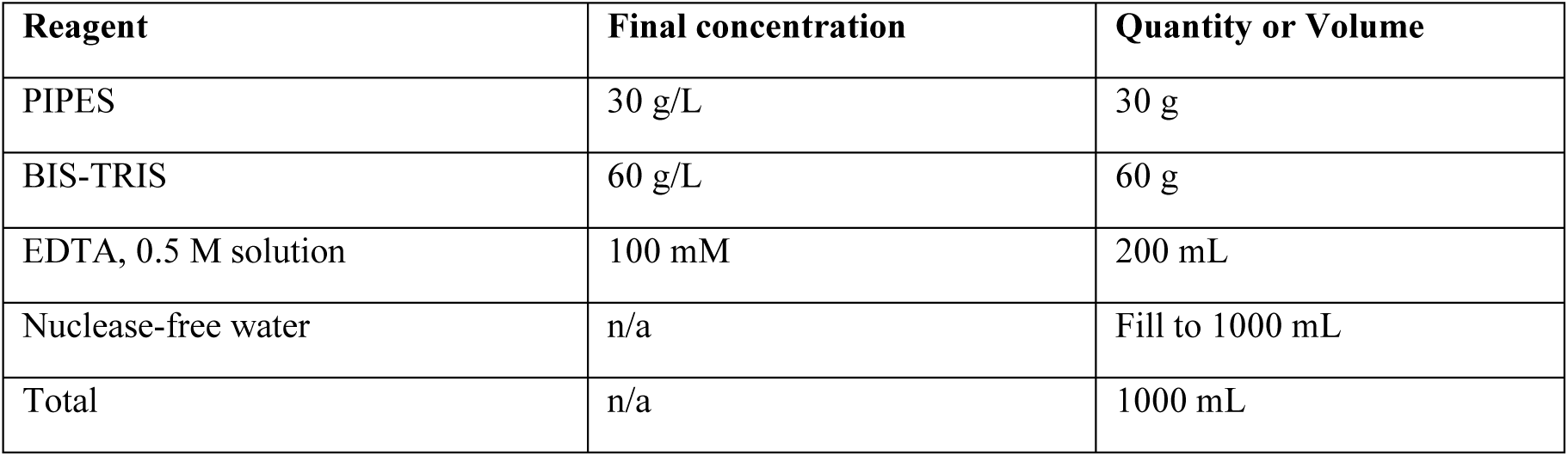
10x BPTE buffer.

| Reagent | Final concentration | Quantity or Volume |
| --- | --- | --- |
| PIPES | 30 g/L | 30 g |
| BIS-TRIS | 60 g/L | 60 g |
| EDTA, 0.5 M solution | 100 mM | 200 mL |
| Nuclease-free water | n/a | Fill to 1000 mL |
| Total | n/a | 1000 mL |

**Table 9.** 2x RNA loading dye.

| Reagent | Final concentration | Quantity or Volume |
| --- | --- | --- |
| Formamide, deionized | 70 % [v/v] | 7 mL |
| 10x BPTE buffer | 10% % [v/v] | 1 mL |
| Bromophenol blue | 0.025% [w/v] | 2.5 mg |
| Xylene cyanole | 0.025% [w/v] | 2.5 mg |
| Nuclease-free water | n/a | Fill to 10 mL |
| Total | n/a | 10 mL |

**Table 10.** TRIS hydrochloride 500 mM stock solution (pH 6.8). Adjust pH to 6.8.

| Reagent | Final concentration | Quantity or Volume |
| --- | --- | --- |
| TRIS hydrochloride | 500 mM | 78.8 g |
| Nuclease-free water | n/a | Fill to 1000 mL |
| Total | n/a | 1000 mL |

**Table 11.** TRIS hydrochloride 1.5 M stock solution (pH 8.8). Adjust pH to 8.8.

| Reagent | Final concentration | Quantity or Volume |
| --- | --- | --- |
| TRIS hydrochloride | 1.5 M | 236.4 g |
| Nuclease-free water | n/a | Fill to 1000 mL |
| Total | n/a | 1000 mL |

**Table 12.** 10% SDS stock solution.

| Reagent | Final concentration | Quantity or Volume |
| --- | --- | --- |
| SDS ultra-pure | 10% [w/v] | 100 g |
| Nuclease-free water | n/a | Fill to 1000 mL |
| Total | n/a | 1000 mL |

**Table 13.** 2x SDS-PAGE sample loading buffer.

| Reagent | Final concentration | Quantity or Volume |
| --- | --- | --- |
| TRIS hydrochloride 1 M stock solution | 100 mM | 1 mL |
| Glycerol Formal | 20% [v/v] | 2 mL |
| Bromophenol blue | 0.03% [w/v] | 3 mg |
| 10% SDS stock solution | 2% [v/v] | 2 mL |
| Nuclease-free water | n/a | Fill to 10 mL |
| Total | n/a | 10 mL |

**Table 14.** 10% Ammonium peroxydisulphate stock solution.

| Reagent | Final concentration | Quantity or Volume |
| --- | --- | --- |
| Ammonium peroxydisulphate | 10% [w/v] | 100 g |
| Nuclease-free water | n/a | Fill to 1000 mL |
| Total | n/a | 1000 mL |

**Table 15.** Transfer buffer (semi-dry).

| Reagent | Final concentration | Quantity or Volume |
| --- | --- | --- |
| TRIS | 48 mM | 5.8 g |
| Glycine | 39 mM | 2.9 g |
| 10% SDS stock solution | 0.04% | 4 mL |
| Nuclease-free water | n/a | Fill to 800 mL, adjust pH to 9.2 |
| Methanol | 20% [v/v] | 200 mL |
| Total | n/a | 1000 mL |

**Table 16.** Ponceau S staining solution.

| Reagent | Final concentration | Quantity or Volume |
| --- | --- | --- |
| Ponceau S | 0.5% [w/v] | 0.5 g |
| Glacial acetic acid | 5% [v/v] | 5 mL |
| Nuclease-free water | n/a | Fill to 100 mL |
| Total | n/a | 100 mL |

**Table 17.** 10x TBS buffer. Adjust pH to 7.5.

| Reagent | Final concentration | Quantity or Volume |
| --- | --- | --- |
| Sodium Chloride | 1.5 M | 87.7 g |
| TRIS hydrochloride | 100 mM | 15.8 g |
| Nuclease-free water | n/a | Fill to 1000 mL |
| Total | n/a | 1000 mL |

**Table 18.**
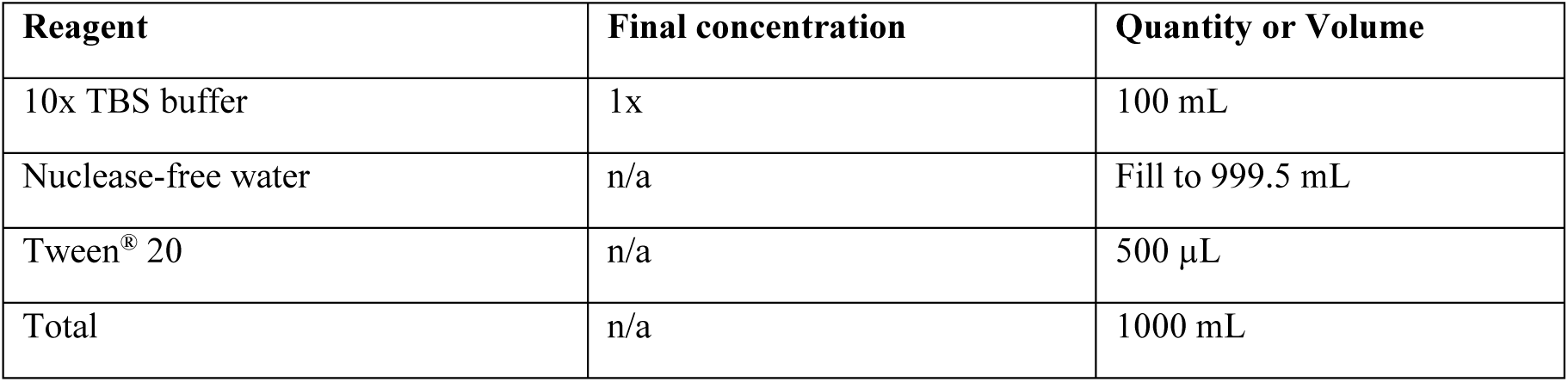
TBS-T buffer.

| Reagent | Final concentration | Quantity or Volume |
| --- | --- | --- |
| 10x TBS buffer | 1x | 100 mL |
| Nuclease-free water | n/a | Fill to 999.5 mL |
| Tween® 20 | n/a | 500 µL |
| Total | n/a | 1000 mL |

**Table 19.**
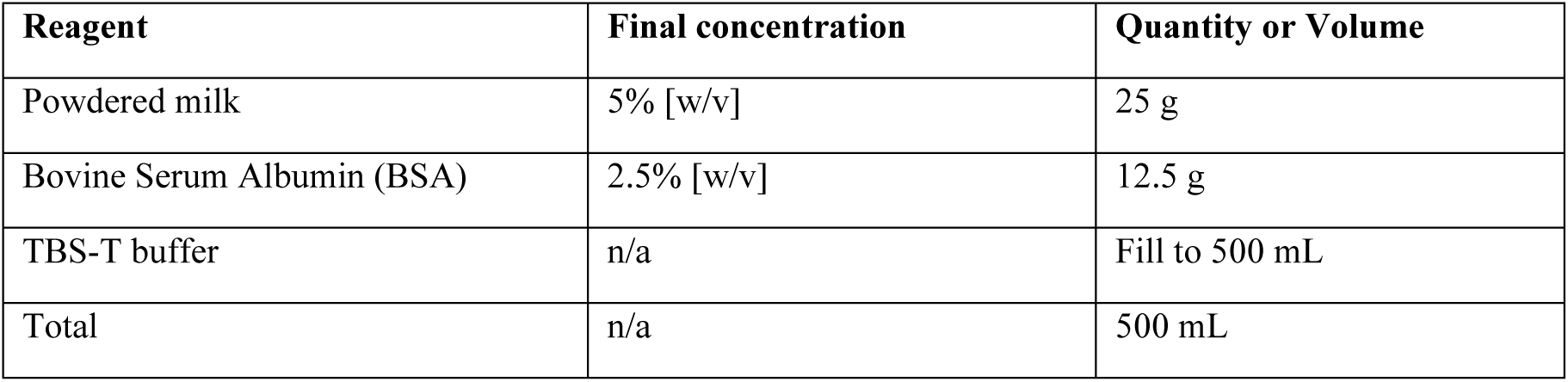
Blocking buffer.

| Reagent | Final concentration | Quantity or Volume |
| --- | --- | --- |
| Powdered milk | 5% [w/v] | 25 g |
| Bovine Serum Albumin (BSA) | 2.5% [w/v] | 12.5 g |
| TBS-T buffer | n/a | Fill to 500 mL |
| Total | n/a | 500 mL |

## Recipes

### Laboratory supplies

1. Amersham^™^ Protran^™^ 0.45 µm NC Nitrocellulose Western Blotting Membrane (Cytiva, catalog number: 10600007)
2. Eppendorf Research^®^ Plus Pipets (Eppendorf, catalog numbers: 3123000012, 3123000020, 3123000039, 3123000055, 3123000063)
3. Nerbe Plus 1.5 mL PP Microcentrifuge Tube with safelock cap (Nerbe Plus, catalog number: 04-212-1000)
4. Nerbe Plus Pipette Tip PP, classic, 100-1000 µL (Nerbe Plus, catalog number: 07-362-2020)
5. Nerbe Plus Pipette Tip PP, classic, 1-200 µL (Nerbe Plus, catalog number: 07-362-2010)
6. Nerbe Plus Pipette Tip PP, premium surface, 0.1-20 µL (Nerbe Plus, catalog number: 07-362-2005)
7. Nunc™ MicroWell™ 96 Wells, Nuclon^TM^ Delta Surface, white, flat bottom (Thermo Fisher Scientific, catalog number: 136101)
8. Qubit^™^ Flex Assay tube strip (Thermo Fisher Scientific, catalog number: Q33252).
9. Whatman Paper (GE Healthcare, catalog number: 3030-917)

### Equipment

1. ChemiDoc Imaging System (Bio-Rad, catalog number: 12003153)
2. Duomax 1030 gyratory shaker (Carl ROTH, catalog number: H836.1)
3. Eppendorf 5417R Refrigerated Centrifuge (Eppendorf, catalog number: 5429000133)
4. Eppendorf ThermoMixer® (Eppendorf, catalog number: 5382000015)
5. Innova® 44R Shaker (New Brunswick, catalog number: M1282-0006)
6. Mini-PROTEAN Tetra Vertical Electrophoresis Cell (Bio-Rad, catalog number: 1658006FC)
7. NanoDrop^™^ One^C^ (Thermo Fisher Scientific, catalog number: 13400519)
8. PowerPac^™^ Universal Power Supply (Bio-Rad, catalog number: 1645070)
9. Qubit^™^ Flex Fluorometer (Thermo Fisher Scientific, catalog number: Q33327)
10. Spark^TM^ Multimode Microplate Reader (Tecan, catalog number: 735-0399)
11. Sub-Cell GT Horizontal Electrophoresis System (Bio-Rad, catalog number: 1704401)
12. Trans-Blot Turbo Transfer System (Bio-Rad, catalog number: 1704150)
13. Vortex-Genie® 2 mixer (Scientific Industries, catalog number: Z258423)

### Software

1. GraphPad Prism (Graphpad Software, version 8.0.1)
2. Qubit^™^ Software (Thermo Fisher Scientific)
3. SnapGene (GSL Biotech LLC, version 8.2)
4. SparkControl (Tecan, version 2.1)

## Procedure

### A. Obtaining IRES sequence

1. Obtain plasmids containing firefly luciferase (FLuc) coding sequence downstream of various plant virus-derived IRES elements from Addgene Repository (addgene.org) (see list of available IRES elements in section *Biological materials*).
2. As a control for 5’- or cap-derived translation obtain plasmid L4821 from Promega.

### B. Propagation and preparation of plasmid DNA

1. Plasmids containing IRES elements will be provided by Addgene Repository as transformed *E. coli* NEB® Stable cells. Directly proceed with step 4 with these strains.
2. Transform NEB® Stable Competent *E. coli* (High Efficiency) (catalog number: C3040) cells with Luciferase T7 Control DNA (catalog number: L4821) by heat shock method. First, thaw 50 µL competent cells in microcentrifuge tube on ice. Once thawed, add 1 µL of plasmid DNA and mix by flicking the tube.
3. Incubate the mixture for 10 minutes on ice, then perform heat shock at 42 °C for 45 seconds using an Eppendorf ThermoMixer^®^ (catalog number: 5382000015). Immediately incubate the mixture on ice for 2 minutes.
4. Streak cells on LB selection plates containing 100 µg/ml ampicillin (CAS number: 69-52-3) and 1.5 % [w/v] of agar (CAS number: 9002-18-0).
5. Select a single colony of each strain of interest and transfer it into 5 ml LB liquid media (see Table 1) containing 100 µg/ml ampicillin for plasmid propagation.
6. Incubate the bacteria overnight, or at least 12 h at 37 °C in an Innova^®^ 44R Shaker (catalog number: M1282-0006) at 250 rpm.
7. Pellet 5 ml of bacterial culture in a 2 ml reaction tube. **Pause point:** Bacteria pellets could be stored at -20 °C for several days. Storage exceeding several weeks will decrease plasmid amounts and sequence qualities.
8. Purify plasmid DNA using Monarch® Plasmid Miniprep Kit (catalog number: T1010) according to manufacturer’s instructions and resuspend DNA in 30 µL of nuclease-free water. Determine DNA concentration using a NanoDrop^™^ One^C^ (catalog number: 13400519). DNA yields should vary between 0.1 - 0.4 µg/µL but should exceed 0.1 µg/µL for downstream procedures. Verify DNA integrity by gel electrophoresis as described in step C. Furthermore, confirm sequence identity during the first execution of the protocol by oxford nanopore sequencing offered by Eurofins genomics (https://eurofinsgenomics.eu/en/custom-dna-sequencing/eurofins-services/whole-plasmid-sequencing) or equivalent sequencing providers.
9. Store plasmid DNA at -20 °C for up to two years or proceed with step D.

### C. DNA gel electrophoresis

1. For confirmation of DNA integrity, prepare a 0.8 % [w/v] TBE agarose gel by boiling 0.8 g of Agarose NEEO Ultra-Quality (catalog number: 2267.4) in 100 mL 0.5x TBE buffer (Table 2) for 3 min in a wide-mouth flask within a microwave at 600 W.
2. Add 5 µL of ROTI^®^GelStain (catalog number: 3865.1) (1:20.000 [v/v]) to your gel solution before pouring. Add a gel comb with enough pockets to load at least the number of samples plus an additional pocket for a sufficient size standard. We recommend using the 1 kb Plus DNA Ladder (catalog number: N3200) (size range: 100 bp to 10 kb).
3. Let the TBE agarose gel solidify for approximately 20-30 min at room temperature (RT) before transferring it into an electrophoresis chamber prepared with 0.5x TBE running buffer.
4. Prepare a size standard and the samples from the transcription reaction with 6x DNA Loading Dye (see Table 4) and transfer the mixture into the pockets. Depending on the equipment, 10-20 µL may be loaded into the gel per lane. We recommend separating around 500 ng of DNA per lane. Dependent on electrophoresis equipment and imager, detection might be possible between 50-1000 ng DNA per lane.
5. Run gel for 45 min at 120 V using a PowerPac^™^ Universal Power Supply (catalog number: 1645070) with a Sub-Cell GT Horizontal Electrophoresis System (Bio-Rad, catalog number: 1704401) and visualize the DNA using the ChemiDoc Imaging System (catalog number: 12003153) using the ethidium bromide setting with “Auto Optimal” exposure time.

### D. Plasmid linearization

1. Plasmids containing the IRES elements have a T7 RNA polymerase promoter region to recruit T7 RNA polymerase. The transcripts contain the IRES sequence upstream the FLuc CDS both flanked by the *Ct*SLP group II intron [14]. The *Ct*SLP sequence does not mediate autocatalytic self-splicing within the plasmids listed above. A restriction enzyme recognition site downstream of the FLuc coding sequence allows plasmid linearization enabling run-off transcription. In addition, superhelicity of non-linearized plasmids limits access of T7 RNA polymerase to the promoter region *in vitro*. Linearization relaxes the plasmid DNA and increases the RNA yield of *in vitro* translation reactions. Therefore, set up the restriction reaction for plasmid linearization of the IRES containing plasmids and the control plasmids (Table 20).
2. Incubate restriction reaction for 30 minutes at 37 °C in an Eppendorf ThermoMixer® (Eppendorf, Cat. No. 5382000015). **Note:** The availability of linearized plasmid linearly affects the amount of synthesizable RNA. Especially, the yield of precipitated plasmid after plasmid linearization could vary drastically between first-time and experienced users. Experienced users typically achieve recovery rates between 50% and 80%, whereas recovery may drop below 30% for novice users. This reaction scale is sufficient for a one-time experiment, if more experiments are planned, the reaction should be scaled up accordingly.
3. Adjust the volume to 200 µL with nuclease-free water and add 20 µL of 3 M sodium acetate (pH 5.2) (see Table 5).
4. Add 200 µL of 2-propanol (CAS number: 67-63-0) and vortex thoroughly.
5. Incubate at least for 30 min at -20°C or overnight before centrifuging at 4°C for 20 min at 14,000 x g in an Eppendorf 5417R Refrigerated Centrifuge (Eppendorf, Cat. No. 5429000133).
6. Carefully discard the supernatant and avoid touching the pellet. Quick-spin the reaction tube again to get rid of residual alcohol.
7. Add 500 µL of ice-cold 70% ethanol (Table 7), centrifuge at 4°C for 5 min at 14,000 x g and discard the supernatant to wash the pellet **Caution:** Do not vortex during the washing steps.
8. Repeat the washing step at least once.
9. Quick-spin at the end to get rid of residual alcohol, air-dry at RT for 5 min and resuspend in 15 µL of nuclease-free-water.
10. Determine DNA concentration using a NanoDrop^™^ One^C^ (catalog number: 13400519). DNA yields should vary between 0.1 - 0.16 µg/µL but should be minimum 0.1 µg/µL for downstream procedures representing at least 50 % of DNA recovery. Confirm DNA integrity and linearization by gel electrophoresis as described in step C. Compare equivalent amounts of DNA before and after linearization.
11. Store linearized DNA at -20 °C for up to two years or proceed with step E.

### E. *In vitro* transcription

1. Synthesize RNA using HiScribe^®^ T7 Quick High Yield RNA Synthesis Kit (catalog number: E2050S). Set up the reaction as described (Table 21). For producing capped transcripts, add 6 µL of 3′-O-Me-m^7^G(5’)ppp(5’)G RNA Cap Structure Analog (catalog number: S1411S) per reaction and reduce the volume of nuclease-free water by the same.
2. Incubate *in vitro* transcription reactions at 37 °C for 2 h in an Eppendorf ThermoMixer® (catalog number: 5382000015). Add 2 µL of DNase I (catalog number: M0303S) and 18 µL of nuclease-free water, vortex shortly and spin down. Incubate additional 20 min at 37 °C.
3. Place reaction on ice to stop transcription. Precipitate RNA by adding 22.7 µL of 8 M lithium chloride (see Table 6) solution, which will result in a final lithium chloride concentration of 2.5 M.
4. Incubate for at least 30 min or overnight at -20 °C. Centrifuge at 4 °C for 30 min at 14.000 xg to pellet the RNA.
5. Discard the supernatant and add 200 µL of 70% ethanol (see Table 7) to wash the pellet. Centrifuge at 4 °C for 5 min at 14,000 x g and discard the supernatant again. **Caution:** Do not vortex during the washing steps.
6. Repeat the washing step at least once.
7. Quick-spin at the end to get rid of residual alcohol, air-dry at room temperature for 15 min and resuspend in 30 µL of nuclease-free-water. **Caution:** Incomplete ethanol removal may inhibit downstream applications. However, over-drying the pellet will make resuspension difficult.
8. Determine RNA concentration using Qubit^™^ Flex Fluorometer (catalog number: Q33327). First, dilute 2 µL of transcription reaction with 18 µL of nuclease-free water. Add 1 µL of the diluted RNA to 199 µL Qubit^™^ RNA Broad Range Assay Kit mixture (catalog number: Q10210). Mix everything in 1.5 ml microcentrifuge tube, transfer the mixture to a Qubit^™^ Flex Assay tube strip (catalog number: Q33252) and measure samples in the Qubit Flex Fluorometer. RNA concentration of your undiluted purified transcription reactions should vary between 2 - 4 µg/µL but should be minimum 1.4 µg/µL for downstream applications.
9. Store RNA at -20 °C for short (up to two weeks) or at -80 °C for longer durations. Otherwise confirm transcription homogeneity by gel electrophoresis described in step F.

### F. RNA gel electrophoresis

1. To confirm transcript homogeneity, prepare a 0.8 % [w/v] BPTE agarose gel by boiling 0.8 g of Agarose NEEO Ultra-Quality (catalog number: 2267.4) in 100 mL 1x BPTE buffer (prepared from 10x BPTE buffer; Table 8) for 3 min in a wide-mouth flask within a microwave at 600 W.
2. Add 5 µL of ROTI^®^GelStain (catalog number: 3865.1) (1:20.000 [v/v]) to your gel solution before pouring. Add a gel comb with enough pockets to load at least the number of samples plus an additional pocket for a sufficient size standard. We recommend using the RiboRuler High Range RNA Ladder (catalog number: SM1821) (size range: 200 to 6000 nt) or the RiboRuler Low Range RNA Ladder (catalog number: SM1831) (size range: 100 to 1000 nt). The transcript lengths of the provided plasmids are listed in Table 22.
3. Let the BPTE agarose gel cool down for approximately 20-30 min at RT before transferring it into an electrophoresis chamber prepared with 1x BPTE buffer (prepared from 10x BPTE buffer; Table 8).
4. Prepare a size standard and the samples from the transcription reaction with 2x RNA Loading Dye (Table 9), denature samples at 95 °C for 10 min in an Eppendorf ThermoMixer® (catalog number: 5382000015) and place directly on ice, before transferring the mixture to the individual pockets. Depending on the equipment, 10-20 µL may be loaded into the gel per lane. We recommend separating around 500 ng of RNA per lane. Detection, dependent on electrophoresis equipment and imager, might be possible between 50-1000 ng per lane.
5. Run gel for 45 min at 120 V using a PowerPac^™^ Universal Power Supply (catalog number: 1645070) with a Sub-Cell GT Horizontal Electrophoresis System (Bio-Rad, catalog number: 1704401), and visualize the RNA using the ChemiDoc Imaging System (catalog number: 12003153) using the ethidium bromide setting with auto optimal exposure time. Transcripts should show a single and homogenous band as shown in Figure 1. Smearing below the expected band indicates degradation by RNase contamination.

### G. *In vitro* translation

1. Thaw Wheat Germ Extract (catalog number: L418A) and Amino Acid Mixture, Complete (L4461) on ice. **Caution:** Avoid more than two freeze-thaw cycles of Wheat Germ Extract to preserve its activity.
2. Prepare the following reaction mix (Table 23) on ice with at least two replicates. Use 5 µg of the control RNA per replicate and equimolar amounts of IRES transcripts in relation to the control. Use the NEBioCalculator tool (https://nebiocalculator.neb.com) to accurately calculate molar masses of your input transcripts. Otherwise, reporter activity will not be comparable between the individual transcripts, resulting in unclear IRES efficiency. To calculate the appropriate RNA amounts from provided constructs, refer to Table 22, which details the length of each transcript. For qualitative experiments, use 5 µg per transcript and reaction.
3. Incubate the translation reaction at 25 °C for 2 h in an Eppendorf ThermoMixer® (Eppendorf, Cat. No. 5382000015).
4. Place the translation reaction on ice until you proceed with H or store it at -20 °C up to one week. **Caution:** Signal intensity in reporter assays after freeze-thaw cycles may decrease.

### H. Reporter activity measurement

1. Prepare a 1:10 dilution by combining 2.5 µL translation reaction with 22.5 µL nuclease-free water. **Critica**l: Measuring the non-diluted translation reaction can result in very low signal.
2. Equilibrate reconstituted ONE-Glo™ Reagent (catalog number: E6110) to room temperature.
3. Dispense 25 µL diluted translation reaction into a well of a white, flat-bottom Nunc™ MicroWell™ 96-well Nuclon^TM^ Delta Surface microwell plate (catalog number: 136101) and add 25 µL ONE-Glo™ reagent to each well. Perform at least two technical replicates for each biological replicate.
4. Transfer plate into a Spark^TM^ Multimode Microplate Reader (Tecan) and perform a 3 mm orbital shake for 30 s at 180 rpm before incubating the reaction at 25 °C for 3 min.
5. Measure luminescence of each well with 1000 ms integration time per well.
6. A successful translation reaction produces a stable luminescence signal in samples containing IRES-luciferase RNA, while control reactions lacking RNA show no signal. Comparative luminescence measurements allow rank ordering of translation efficiency of candidate IRES sequences (Figure 2). Modifications to mRNAs intended to enhance translation can also be assessed in this manner.

### I. Western blot analysis

1. Perform Western blot analysis for a secondary verification of the translation process. The transcripts from the provided constructs (see Table 22) translate a 3xHA tag at the C-terminus of the firefly luciferase. Mix 2.5 µL of the translation reaction with 7.5 µL nuclease-free water and 10 µL 2x SDS-PAGE sample loading buffer (see Table 13). Denature in Eppendorf ThermoMixer® (catalog number: 5382000015) at 65° C for 15 minutes.
2. Prepare separating and stacking gels for SDS-PAGE according to Table 24 and Table 25, respectively. Pour separating gel solution into cast while leaving 2 cm empty for the stacking gel. Cover the separating gel with 70% ethanol and let solidify for 20 minutes or until the remainder of the separating gel solution has fully polymerized. Remove the entire ethanol and pour stacking gel solution on top of solidified separating gel, insert comb with a suitable number of wells. Wait 20 minutes or until the remainder of the stacking gel solution has fully polymerized.
3. Transfer fully hardened polyacrylamide gel into Mini-PROTEAN Tetra Vertical Electrophoresis Cell (catalog number: 1658006FC) and load with 17 µL of prepared protein sample. Load 5 µL of a prestained protein ladder such as the Precision Plus Protein™ Dual Xtra Prestained Protein Standards (catalog number: 1610377) (size range: 2 to 250 kDa). Perform electrophoresis for 20 minutes at 80 V and followed by 55 minutes at 150 V using the PowerPac^™^ Universal Power Supply (catalog number: 1645070).
4. Remove the polyacrylamide gel from the chamber and carefully separate from the cast. Prepare the transfer sandwich in the tray of the Trans-Blot Turbo Transfer System (catalog number: 1704150). First, stack three Whatman papers (catalog number: 3030-917) (8.5 x 7.3 cm) previously soaked in Transfer buffer (semi-dry) (see Table 15). Activate the nitrocellulose membrane (8.5 x 7.3 cm) (catalog number: 10600007) by soaking it in Transfer buffer (semi dry) for one minute, then place it on the Whatman papers. Cut a corner to designate it as the bottom right corner of the membrane for orientation in later steps. Finish transfer sandwich by adding three more soaked Whatman papers on top. Blot by using the StandardSD program (25 V and 1 A for 30 min) for a Mini gel in the Trans-Blot Turbo Transfer System. **Critical:** Avoid letting the nitrocellulose membrane dry and take care to remove all bubbles during transfer sandwich assembly.
5. Confirm protein transfer by Ponceau S staining. First, stain the membrane with 10 mL Ponceau S staining solution (see Table 16) for 2 minutes and wash with deionized water for 2 minutes with gentle agitation on a Duomax 1030 gyratory shaker (catalog number: H836.1) to remove background staining. Document the stained membrane using the ChemiDoc Imaging System and remove the remaining stain by washing with deionized water. Block the membrane by adding 20 mL of blocking buffer (see Table 19) and incubate for 1 hour at room temperature with gentle agitation on a gyratory shaker. **Pause point:** Blocking can also be performed overnight at 4°C with gentle agitation on a gyratory shaker.
6. Wash the membrane three times with 20 mL TBS-T (see Table 18) by incubating on gyratory shaker. Dilute 2 µL of the HA Tag Recombinant monoclonal antibody (catalog number: 81290-1-RR) in 15 mL Blocking buffer and incubate for 2 h with gentle agitation on a gyratory shaker. **Pause point:** Incubating the membrane with the primary antibody can also be performed overnight at 4° C with gentle agitation on a gyratory shaker. This can increase signal intensity in some cases.
7. Wash the membrane with 20 mL of TBS-T on a gyratory shaker, incubating for 5 minutes per washing step. Add 1 µL of Peroxidase IgG Fraction Monoclonal Mouse Anti-Rabbit IgG antibody (catalog number: 211-032-171) diluted in 15 mL blocking buffer and incubate for 2 h at room temperature with gentle agitation on a gyratory shaker. Remove the excess secondary antibody by washing the membrane three times with 20 mL of TBS-T for 5 minutes each.
8. Generate chemiluminescence signal by adding 7 mL of Clarity Western ECL Substrate on top of the membrane (catalog number: 1705061) and incubate for 2 minutes. Document the blot in the ChemiDoc Imaging System using the multichannel setting with a colorimetric and chemiluminescence channel. Illuminate using the Auto-Optimal setting. Blots should show distinct signal at ∼65 kDa (see Figure 3).

## Data analysis

Extract raw luminescence data in counts per second (CPS) from SparkControl output Excel file. Normalize each measurement to desired control RNA by dividing by the mean CPS of the control RNA replicates. Use GraphPad Prism (version 8.0.1) to perform statistical analysis, conduct a one-way ANOVA to compare each IRES-construct group with the control RNA group. Create graphical representations (see bar plot in Figure 2) in GraphPad Prism. Verify the resulting data by confirming protein expression in Western blot analyses (see Figure 3).

**Figure 2.**
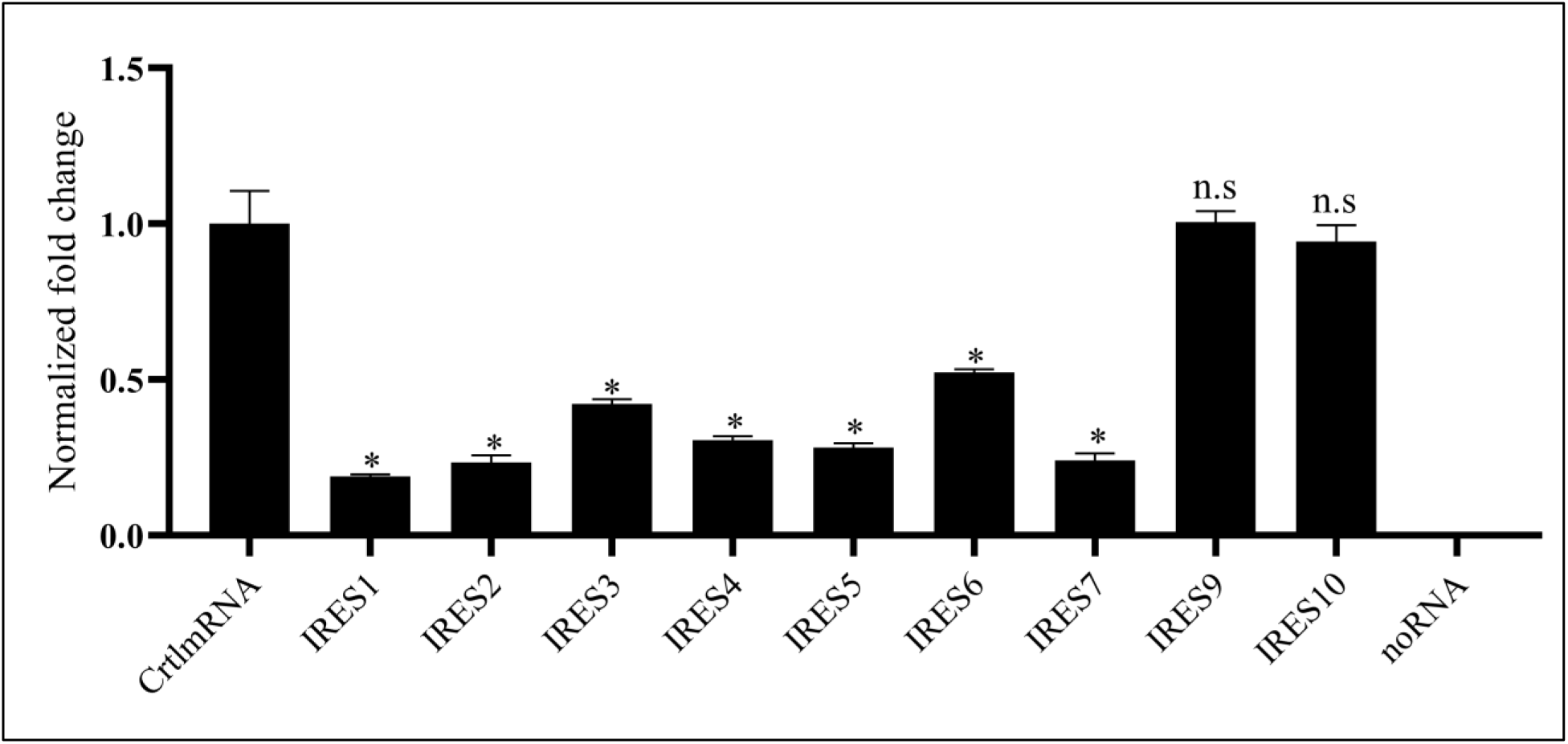
Luminescence measurements from IRES-luciferase constructs normalized to CtrlmRNA. Luminescence readouts were normalized to CtrlmRNA. Statistical significance is indicated by * (p < 0.0001), bars labeled “n.s” are not significantly different.

**Figure 3.**
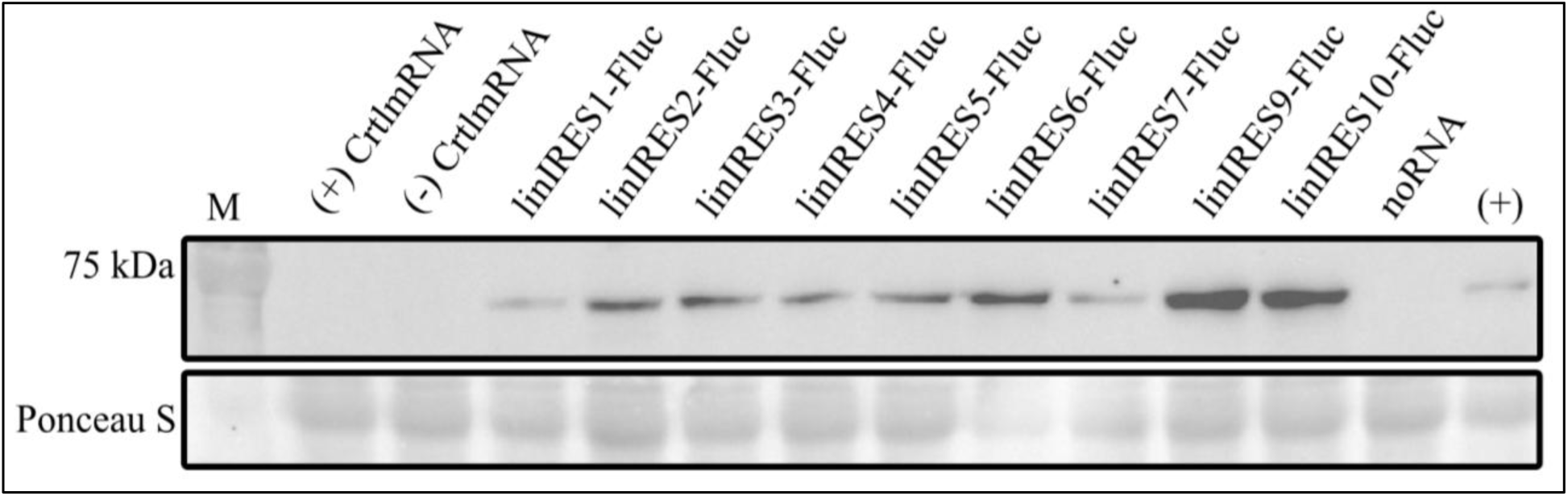
Western blot analysis of the *in vitro* translation reaction. The “(+) CtrlmRNA” and” (-) CtrlmRNA” lanes correspond to the RNA transcribed from the L4821 plasmid, with (+) possessing a 3′-O-Me-m7G(5’)ppp(5’)G RNA Cap Structure Analog and (-) possessing no cap structure. These lanes show no signal due to the control plasmid having not 3xHA tag. The remaining lanes show a distinct signal at ∼65 kDa (FLuc (∼61 kDa) fused with 3xHA tag (∼4.4 kDa)) confirming the expression of FLuc. The lane “noRNA” contains the negative control with no RNA added to the translation reaction. The lane “(+) “contains a 3xHA-tagged FLuc as a positive control.

**Table 20.** Plasmid linearization reaction mix.

| Reagent | Final concentration | Quantity or Volume |
| --- | --- | --- |
| Plasmid | 0.1 μg/μL | 3 μg |
| rCutsmart™ Buffer | 1x | 3 μL |
| IRES construct: DraIII-HF® | 5 u/μL | 0.5 μL |
| L4821: AfeI | | 1 μL |
| Nuclease-free water | n/a | Fill to 30 μL |
| Total | n/a | 30 μL |

**Table 21.** *In vitro* transcription reaction mix. All reagents are included in the kit.

| Reagent | Final concentration | Quantity or Volume |
| --- | --- | --- |
| NTP Buffer Mix | 10 mM each NTP | 15 μL |
| Template DNA | 1 μg | n/a |
| DTT (0.1 M) | 5 mM | 1.5 μL |
| T7 RNA Polymerase Mix | n/a | 3 μL |
| Nuclease free water | n/a | Fill to 30 μL |
| Total | n/a | 30 μL |

**Table 22.**
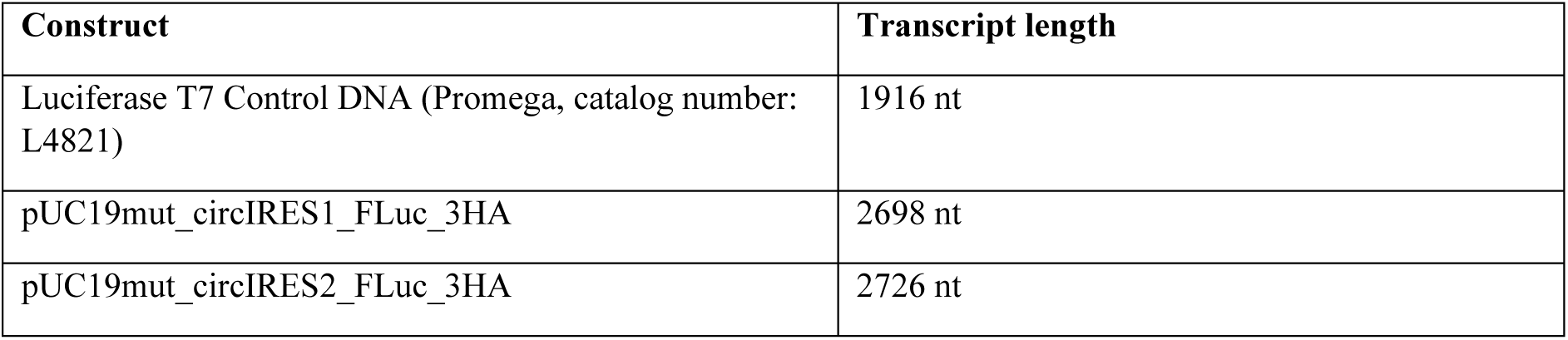
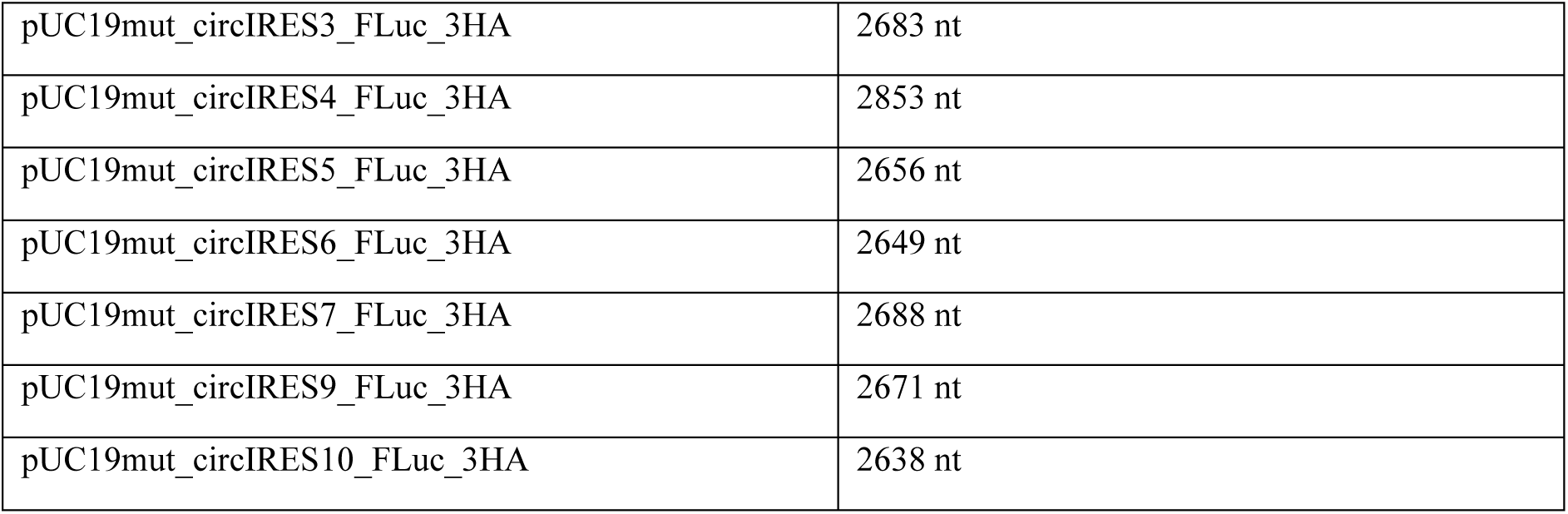
Transcripts and their respective lengths.

**Table 23.**
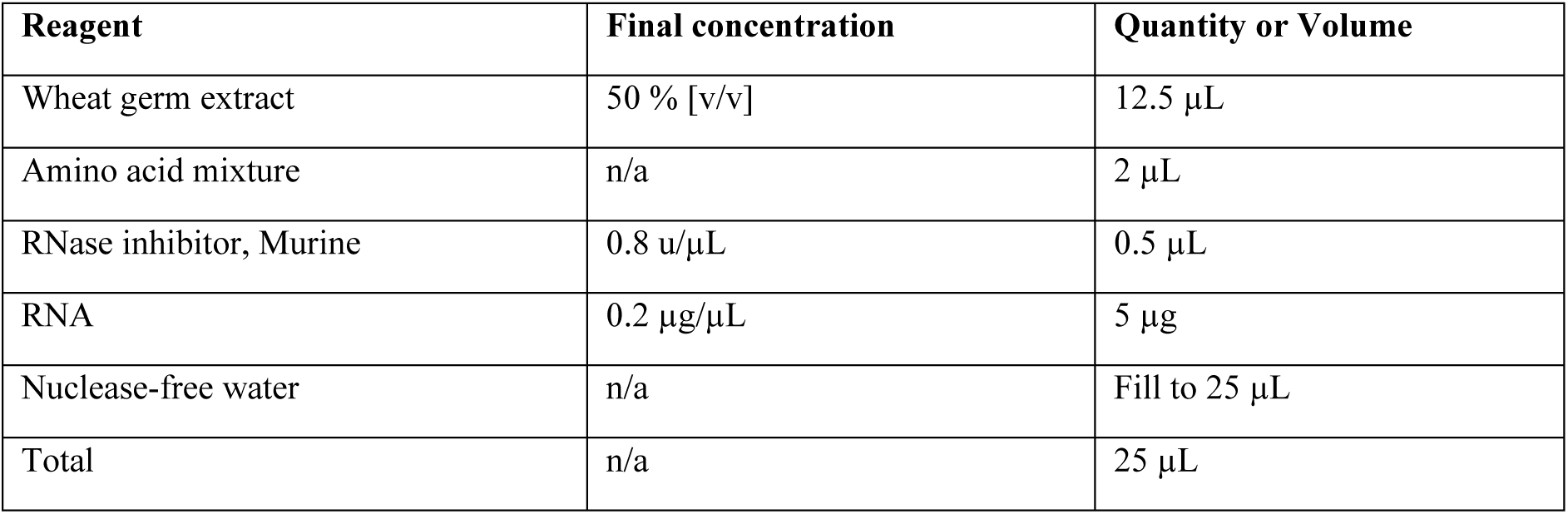
*In vitro* translation reaction mix.

**Table 24.** Recipe for the 12% SDS-PAGE separating gel.

| Reagent | Final concentration | Quantity or Volume |
| --- | --- | --- |
| ROTIPHORESE® Gel 40 (37.5:1) | 12% | 2.4 mL |
| TRIS hydrochloride 1.5 M stock solution (pH 8.8) (Table 11) | 375 mM | 2 mL |
| 10% SDS stock solution (Table 12) | 0.1% | 80 µL |
| 10% Ammonium peroxydisulphate stock solution (Table 14) (Add when ready to pour) | 0.1% | 80 µL |
| TEMED (Pour immediately after adding) | n/a | 8 µL |
| Nuclease free water | n/a | 3.46 mL |
| Total | n/a | 8 mL |

**Table 25.** Recipe for the 6% SDS-PAGE stacking gel.

| Reagent | Final concentration | Quantity or Volume |
| --- | --- | --- |
| ROTIPHORESE® Gel 40 (37.5:1) | 6% | 0.75 mL |
| TRIS hydrochloride 500 mM stock solution (pH 6.8) (Table 10) | 125 mM | 1.25 mL |
| 10% SDS stock solution (Table 12) | 0.1% | 50 µL |
| 10% Ammonium peroxydisulphate stock solution (Table 14) (Add when ready to pour) | 0.1% | 50 µL |
| TEMED (Pour immediately after adding) | 0.1% [v/v] | 5 µL |
| Nuclease free water | n/a | 2.9 mL |
| Total | n/a | 5 mL |

## Validation of protocol

Validation information supporting this protocol is detailed within the procedure section. Figure 1 shows the validation of the *in vitro* transcription step. This validation step should be performed before each *in vitro* translation to ensure RNA integrity. Figure 2 shows luminescence measurements comparing different IRES-constructs to a control mRNA. This data was obtained from an experiment containing biological triplicates and technical duplicates, resulting in 6 measurements per construct. The measurements yielded CPS ranging from 9,000 to 400,000, while the negative controls consistently measured below 200 CPS. Figure 3 shows a secondary validation of the *in vitro* translation reaction using Western blot analysis. Ponceau S staining was used as a loading control.

## General notes and troubleshooting

### General notes

1. Keep all reagents RNase-free and sterile to prevent RNA degradation.
2. Perform all verification steps to ease troubleshooting.
3. Use high quality linearized plasmid templates to ensure consistent RNA yield.
4. Perform assay in appropriate number of biological replicates.
5. Incorporate reliable controls.
6. Wear gloves when handling samples.

## Troubleshooting

Problem 1: Low luminescence signal.

Possible cause: RNA degradation or too high WGE concentration when taking measurement.

Solution: Verify RNA integrity with post-translation Northern blot analysis and avoid RNase contamination. Perform serial dilution of translation reaction before measuring.

Problem 2: Unspecific signal in Western blot analysis. Possible cause: Too much translation reaction loaded.

Solution: Determine optimal amount of translation reaction by loading different dilutions.

## Acknowledgments

Conceptualization, M.C. and T.Sc.; Investigation, M.C. Writing—Original Draft, M.C.; Writing—Review & Editing, M.C., T.St., A.K. and T.Sc.; Funding acquisition, T.S.; Supervision, T.S.; This work was funded by the German Federal Ministry of Research, Technology and Space, project number 031B1588. We thank Prof. Dr. Stefan Schneuwly for providing access to the Spark^™^ Multimode Microplate Reader by Tecan. We thank Maria Laura Mihaila and Maximilian Klein for creating the pUC19mut backbone, amplifying the IRES sequences and cloning precursor plasmids which directly lead to the creation of the here provided constructs.

